# Understandable Multifunctionality Measures Using Hill Numbers

**DOI:** 10.1101/2022.03.17.484802

**Authors:** Jarrett E. K. Byrnes, Fabian Roger, Robert Bagchi

**Affiliations:** Department of Biology, University of Massachusetts Boston, Boston, Massachusetts 02125 United States of America; Department of Environmental Systems Science, Institute of Biogeochemistry and Pollutant Dynamics (IBP), ETH Zurich; Department of Ecology and Evolutionary Biology, University of Connecticut, 75 N. Eagleville Road, Storrs, Connecticut, 06269-3043 United States of America

**Keywords:** multifunctionality, ecosystem function, Hill numbers, diversity

## Abstract

In ecology, multifunctionality measures the simultaneous performance of multiple ecosystem functions. If species diversity describes the variety of species that together build the ecosystem, multifunctionality attempts to describe the variety of functions these species perform. A range of methods have been proposed to quantify multifunctionality, successively attempting to alleviate problems that have been identified with the previous methods. This has led to a proliferation of more-or-less closely related metrics which, however, lack an overarching theoretical framework. Here we borrow from the comprehensive framework of species diversity to derive a new metric of multifunctionality. Analogously to the effective number of species used to quantify species diversity, the metric we propose is influenced both by the number of functions as well as, crucially, the evenness of performance levels across functions. In addition, the effective multifunctionality also considers the average level at which the functions are performed. The result is a measure of the cumulative performance of the system were all functions provided equally. The framework allows for the inclusion of the correlation structure among functions, thus allowing it to account for non-independence between functions. We show that the average metric is a special case of the newly proposed metric when all functions are uncorrelated and performed at equal levels. We hope that by providing a new metric of multifunctionality anchored in the rigorous framework of species diversity based on effective numbers, we will overcome the considerable skepticism that the larger community of ecologists has built against indices of multifunctionality. We thereby hope to help popularize this important concept which, as biological diversity, describes a fundamental property of ecosystems and thus lies at the heart of ecology.

## Introduction

The past decade has witnessed the growth of the concept of ecosystem *multifunctionality*, where multifunctionality is defined as a measure of the simultaneous performance of multiple functions. Having existed as a concept in the ecosystem service literature for a long time (Hölting et al 2019) it has arisen, largely independently, in the biodiversity ecosystem function literature. From there, the concept has spread to community ecology writ large (Angelini et al. 2015), invasion biology (Ramus et al. 2017), land management (Nelson et al. 2009, Brandt et al. 2014, Binder et al. 2018), and more. The concept of multifunctionality is broad and can even be applied outside of community and ecosystem ecology - or even outside of ecology altogether. Its adoption as a unifying concept, however, has been slow - and for good reason.

Despite the popularity that the concept gained since it was first explicitly proposed in community ecology over a decade ago (Hector and Bagchi 2007, Gamfeldt et al. 2008), there is still no consensus on how multifunctionality should be quantified. A range of methods have been proposed where each subsequent method attempted to alleviate problems the authors had identified with the previous method, leading to a proliferation of metrics which cannot easily be compared and can lead to inconsistent interpretations of the results (Byrnes et al. 2014a, Gamfeldt and Roger 2017). We do note that this is a difficult problem. Throwing many different ecologists against this problem in a stimulating working group over multiple years (we are thankful for NCEAS for providing this venue) has likely resulted in more ink impregnated on foreheads hitting whiteboards than should be typical, to say nothing of others who have attempted to cut this Gordian knot before and since (Hector and Bagchi 2007, Brandt et al. 2014, Dooley et al. 2015, Rodríguez-Loinaz et al. 2015, Stürck and Verburg 2017, Manning et al. 2018, Meyer et al. 2018, Hölting et al. 2019). In part, the problem is due to a lack of an underpinning body of theory describing how and where multifunctionality should arise. Regardless, this plurality of measurements has hampered a general understanding of multifunctionality and possibly its adoption outside of the subfield of Biodiversity and Ecosystem Function.

Species diversity, for a long time, suffered the same problems that multifunctionality suffers today: there are an uncountable number of metrics to estimate alpha diversity, reaching from the simple count of observed species without regard to their abundance (Richness), wide array of metrics incorporating the relative abundance (Shannon entropy, Simpson index or the Berger-Parker Dominance index to name some of the most common ones), to metrics based on the histograms of the abundance distribution (e.g. Fisher’s alpha) (Magurran and McGill 2010). Yet, starting with MacArthur (1965), then Hill (1973), followed by Jost (2006), and most recently Chao et al. (2014a, 2019), a common framework for species diversity has been developed based on information theory. This framework, the effective number of species, encompasses the vast majority of previous metrics and is able to handle a wide variety of different issues in diversity estimation. We propose it can do the same for multifunctionality.

### The current state of multifunctional affairs

The definition of multifunctionality, the simultaneous performance of multiple functions, (*sensu* Byrnes et al. 2016) presents a challenge in creating a metric. How do we define a metric that captures both the level of performance of a broad suite of functions as well as the distribution of differences in performance among functions? Researchers have sought to capture this question in four ways, after standardizing functions to similar levels in order to prevent apples-to-oranges comparisons.

First, many have opted for simplicity and taking the average of all functions (e.g., Maestre et al. 2012). This approach is, at first glance, appealing, particularly as it provides a metric that can be put on a y-axis while a predictor is on the x-axis. However, it sacrifices crucial information about the system. An arithmetic mean tells us what level of functioning we would expect were we to sample any one function at random. Consider two plots - one where all functions are similar and performing at half their value and one where half of the functions are at their maximum while half are absent. The averaging approach says that they are identical. Geometric averaging (e.g., Hensel and Silliman 2013) appears to get around this problem to some degree, as the geometric mean is less than the arithmetic mean by a function of the variance of the observations (the two means are the same when all observations are the identical). However, given its formulation, one critically low function can dominate the measurement - e.g., if even one function is 0, the geometric mean will be 0.

Second, we have metrics of the Multivariate Diversity Interactions framework (Dooley et al. 2015). This elegant framework allows us to tease apart the importance of correlations between functions and the contribution of different drivers to simultaneous change in those functions. It does not, however, provide a holistic metric of multifunctionality *per se*, much like the overlap approach before it (Hector and Bagchi 2007). This approach from Hector and Bagchi is a generalization of Sørenson-Dice overlap (Dice 1945, Sørensen 1948). It provides key information on redundancy versus unique contributions of species. While it gives us rich information about a system, it lacks holistic interpretability.

Last, we have the threshold approach (Gamfeldt et al. 2008). This approach defines the “presence” of a function if it meets a given threshold of performance, and then sums up the number of functions performing at or above that threshold. There are clear issues with the arbitrariness of thresholds in the absence of relevant theory or management goals. To remedy this, the multiple threshold approach (Byrnes et al. 2014a) seeks to balance the goals of measuring the simultaneous performance of multiple ecosystem functions with the arbitrariness of choosing a threshold of relevance for those functions. The results, however, can be difficult to interpret. Looking at multiple relationships between diversity and threshold-based multifunctionality does not provide a metric *per se*. While there are key metrics that can be extracted from looking at change in slope over different thresholds, the meaning of these quantities in the context of multifunctionality are not obvious. The values can vary dramatically depending on the exact threshold chosen and on the method used to standardize the functions – even for simulated communities where all functions perform identically (Gamfeldt and Roger 2017, Figure S1). In the absence of suitable null-models, and even if the approach yields rich information about multifunctionality *sensu stricto*, it is unwieldy for most if not all who choose to use it, and we have noticed that many of those who do, often report a single threshold in their main text and leave further exploration to supplementary materials anyway. Related, more recent, efforts have sought to use dimensionality-reducing techniques which have yielded metrics that, while useful, similarly lack easily interpretable meaning (Meyer et al. 2018).

### Hill numbers and effective diversity

Over the last 70 years, ecologists studying how to measure species diversity have shown that the vast majority of previous diversity metrics can be organized into a common framework (Macarthur 1965, Hill 1973, Jost 2006, Chao et al. 2014a, 2019). This is true for indices such as Shannon entropy, all Simpson measures, all Renyi entropies, all HCDT or “Tsallis” entropies and species richness (Jost, 2006) and more. All can be expressed as generalized entropies that can be converted to an effective number of species of “order” *q* which specifies the weighting of proportional abundances. The general formula for the diversity of order *q* for *S* species is the following:

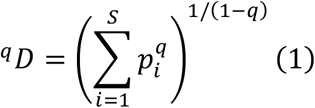

Here, *p*_*i*_ is the relative abundance of the *i*^th^ species and *q* is the weight given to the species’ relative abundances. Species richness, the effective number of species based on Shannon entropy, the effective number of species based on the Simpson index, and the Berger-Parker dominance index are all effective numbers of species of order = 0, 1, 2 and ∞, respectively. (Note that the formula is undefined for *q* = 1, but its limit *q* → 1 is exp(-Σ*p*_*i*_ log *p*_*i*_)). The effective number of species of order *q* is also often referred to as Hill numbers.

We propose leveraging this framework for a more meaningful and less *ad hoc* metric of multifunctionality composed of two parts: (1) the effective *number* of functions that are performed and (2) the *arithmetic mean performance* of the functions that are measured. Our proposed multifunctionality index is then the product of both terms. This approach draws on ideas already swirling in the multifunctionality literature (Brandt et al. 2014, Rodríguez-Loinaz et al. 2015, Stürck and Verburg 2017, Hölting et al. 2019). With this approach, we aim to provide a unifying framework for the measurement of multifunctionality.

### Multifunctionality as the product of effective number of functions and average level of functions

To define the effective number of functions, we begin with a set of measurements on *k* functions (Table 1) that have been standardized to a common scale (i.e., between 0 and 1 where 0 means no function and 1 means maximum level of function). Let *F*_*i*_, *i* ∈ 1, 2, … *K* show the level of function for function *i* (Table 1). The relative proportion a function contributes to the whole is defined as

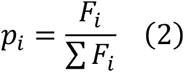

We can now substitute the relative proportion into the formula for the effective number of types given in Eq 1

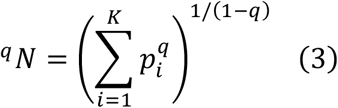

where ^*q*^*N* is the **effective number of functions** for some order *q* (Table 1). *The effective number of functions here translates to the equivalent number of functions were all functions provided at the same level*. Effective number of functions tells us nothing about total level of functioning. Average function can be low or high (see below and Fig. 1). Rather, ^*q*^*N* tells us, given how different functions are performing at different levels, how many functions would we see in an equivalent system where all functions are performing at the same level after weighting for the importance of low-performing functions based on *q*.

**Table 1:**
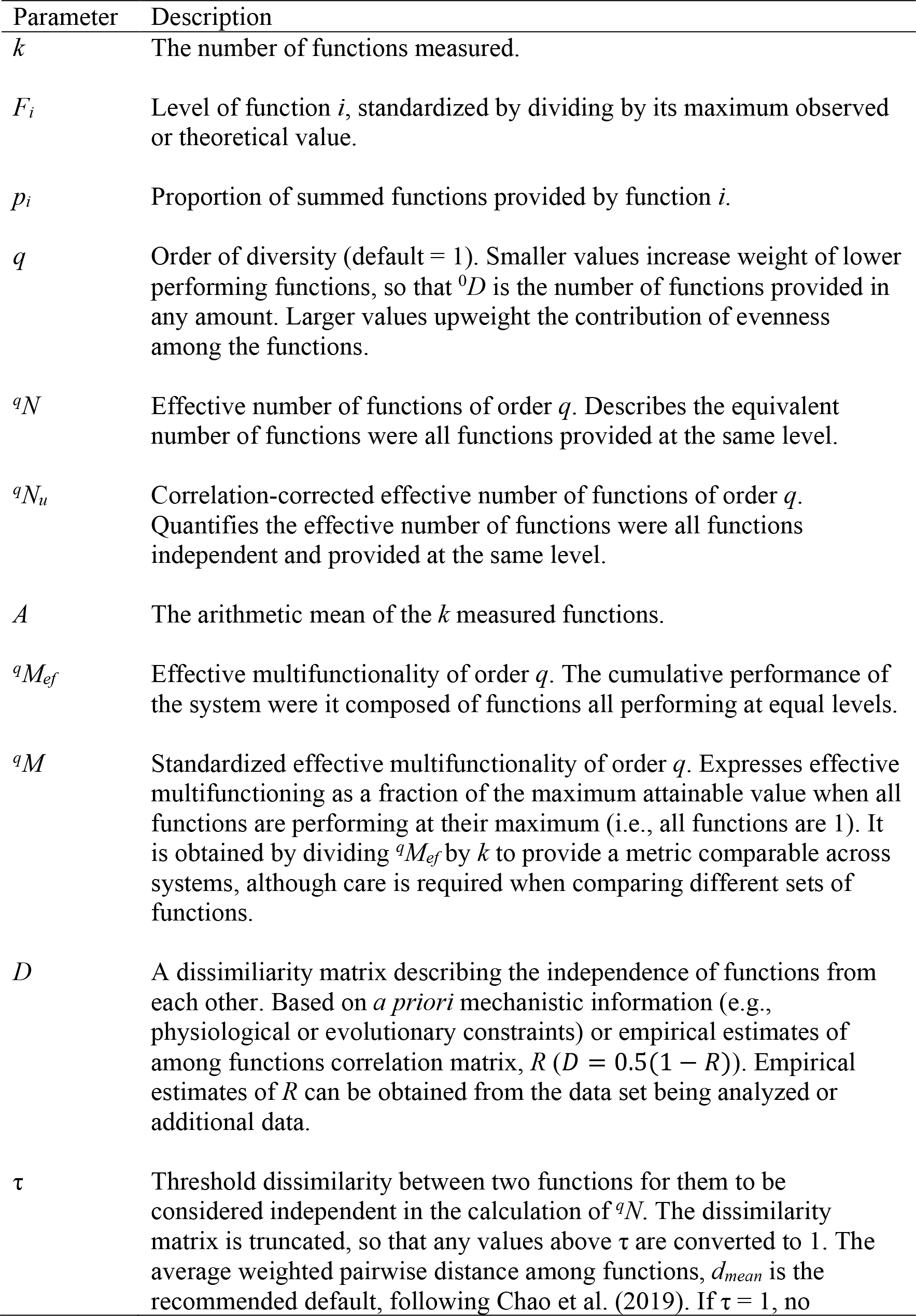
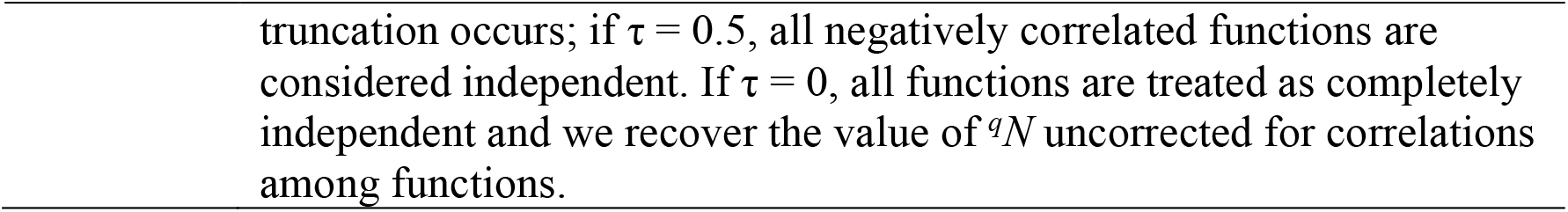
Definitions of parameters used to define the effective multifunctionality of an ecosystem.

**Figure 1.**
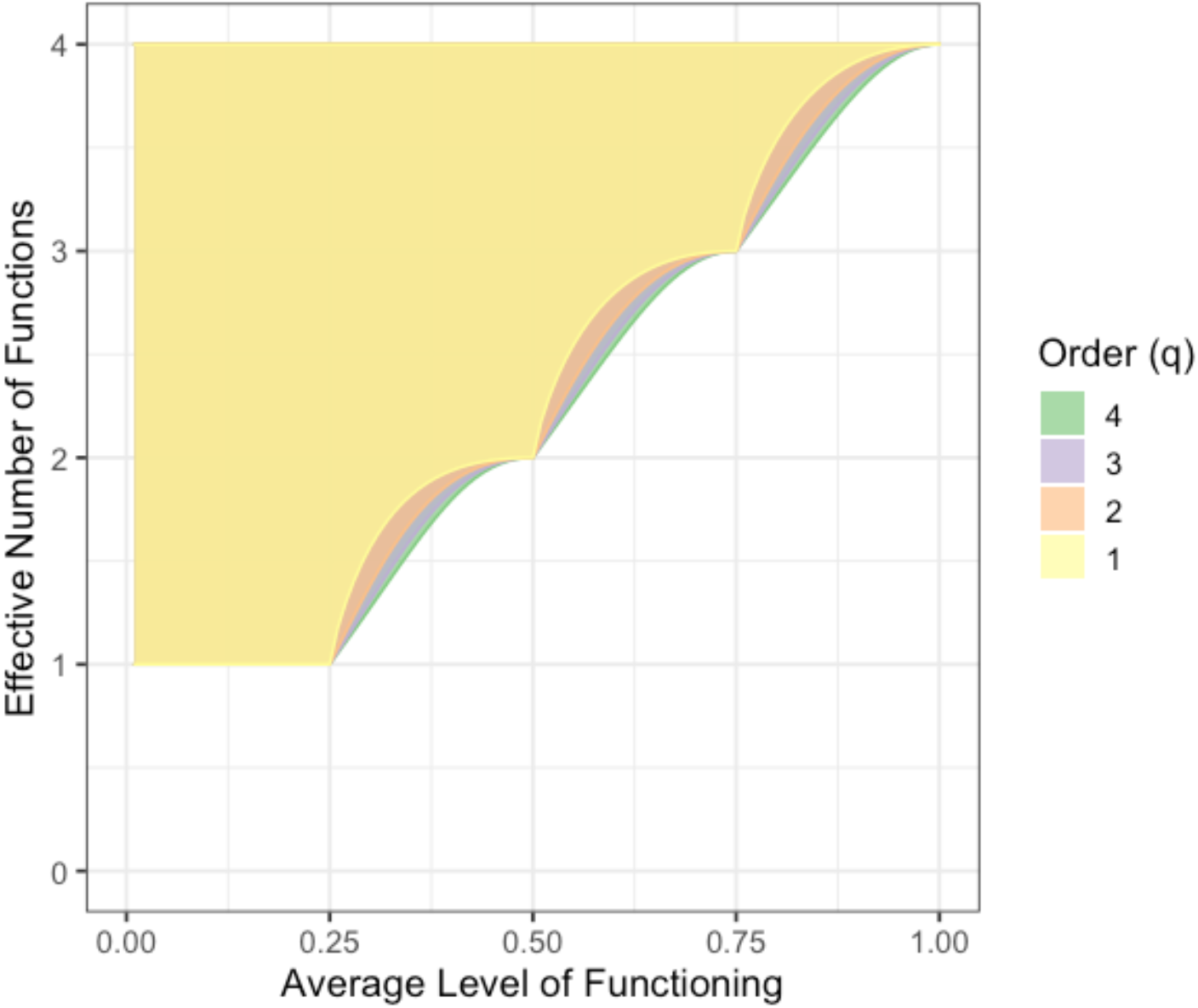
Relationship between effective number of functions, ^*q*^*N* and average level of functions, *A*, for 4 functions at *q* = 1,2,3, and 4. Filled areas show the full range of possible values of effective number of functions, ^*q*^*N*, under different values of average level of function, *A*. Note, areas with a higher order (*q*) do overlap completely with lower orders, save for the additional edge that is visible.

Adapting from Jost (2006), if *q* = 0, this is the total number of functions *K*, which is unimportant as it is just the number of functions measured. In essence, all functions have equal weight regardless of performance. For *q* = 1, the approximation of this function is equivalent to results from Shannon diversity for species shown earlier – which we note has been used as a multifunctionality metric previously (Stürck and Verburg 2017). For *q* > 1, functions performing at higher levels are given greater weight. At *q* = 2, we get results that are equivalent to the number of functions calculated from Simpson’s diversity (inverse Simpson index). If one of our goals is to up-weight high performing functions, *q* = 2 is a reasonable choice, while *q* = 1 is sufficient as it accommodates information about unequal levels of functioning proportional to the relative functional performance. Lower values of *q* would upweight low-performing functions - which might be desirable in certain contexts. The ability to modulate the sensitivity of the metric to high or low performing functions thus provides a strong tool for both ecologists and managers. In the absence of a justification for a particular value of *q*, exploring the robustness of results to different choices of *q* could prove fruitful as it has in biodiversity research. Further exploration of how *q* relates to management goals or ecological theory of multifunctionality would be a fruitful avenue for future research.

Effective number of functions does not accommodate knowledge about the absolute level of functioning in a system - a true metric of multifunctionality. As long as the distribution of *p*_*i*_ is the same, a system with high average function and low average function will look the same. Indeed, for some values of *q* (e.g., *q* = 1), under some scenarios if one function goes up, the effective number of functions can actually drop. To achieve the translation to a metric of multifunctionality, we need to take into account the level at which the functions are performed: the arithmetic mean of the function values standardized to a common scale, which we define as *A* (Table 1). As we are using standardized values as before, *A* will range from 0 to 1.

We can then calculate **effective multifunctionality** of order *q* (Table 1) as the product of both terms. We remind readers that *A* is an expected value – it provides information on the expected level of one function sampled at random from the cluster of functions. Scaling *A* by ^*q*^*N* gives a metric of multifunction summed across the suite of functions – *the cumulative performance of the system were it composed of functions all performing at equal levels*

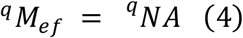

This metric, where ^*q*^*M*_*ef*_ is **effective multifunctionality** for order *q*, will have a maximum value of *K*, the total number of functions measured in the system, as maximum performance is all functions performing at a standardized level of 1. Alternatively, we can standardize by the total number of functions to calculate the fraction of performance achieved by the whole system, of **standardized effective multifunctionality**, ^*q*^*M* (Table 1), which could facilitate future comparisons among studies.

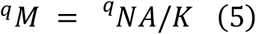

Given that we have scaled by *K*, this metric measures the level of individual functions in an equivalent system where all functions have the same level of performance. However, care has to be taken when comparing multifunctionality values across systems that measure different sets of functions and the value of such comparisons is a topic that eludes consensus even among authors of this article.

Why must we consider the level of function and effective number of functions together in one metric? First, when *A* is less than one, multiple values of ^*q*^*N* are possible depending on the distribution of performance of functions. Second, the average functioning and the effective number of functions are not independent of each other. The upper limit of ^*q*^*N* is *K* by definition. ^*q*^*N* is *K* when all functions are performing at the same level, i.e. *E[A] = F*_*1*_, *F*_*2*_, *F*_*3*_*…F*_*K*_. This can be achieved for any value of *A*, low or high. Thus, ^*q*^*N* has no information about the level of function achieved. The lower limit of ^*q*^*N* (as long as *q* is not 0) occurs at the maximum level of unevenness of *F* across all functions. This lower limit will vary by different levels of *A*. For example, ^*q*^*N* will always be *K* when *A* is 1. Similarly, if all functions but one have a value of 0, then ^*q*^*N* will be 1, as *p*_*i*_ for the non-zero function will be 1. If more than 1 function exceeds 0, then the lower limit is set by a combination of *A* and the number of dominant functions, i.e., the functions performing at *F*_*i*_ = 1 (Figure 1). See Appendix 1 for derivation.

Most importantly, a combined metric satisfies our definition of multifunctionality. High numbers imply both a high level of function and high functional evenness (i.e., p_i_ is close to 1/*K* for all functions). Low values imply that, even if a single function is being optimized, the assemblage of functions as a whole is not performing well. The relationship can also be easily decomposed into its constituent parts for a more detailed examination of its behavior.

Last, from a convenience standpoint, having a single metric allows us to begin to examine it as any other response variable. In the Biodiversity and Ecosystem Functioning world, we might look at additive partitioning in addition to complementary overlap approaches. In global change biology, we can look at the stability, resistance, and resilience of this metric in ecosystems confronting human stressors. This metric can be used just as any other univariate metric in any field, leading to easy adoption of the multifunctionality concept across many fields of endeavor. The options are open.

### A note on standardization

As pointed out in discussions of multifunctionality, how functions are standardized matters (Gamfeldt and Roger 2017, Manning et al. 2018). First is the choice of direction - what implies positive function? When services are being examined explicitly, this is hopefully a straightforward choice - although consider tradeoffs between nutrient cycling rates and storage as a tricky context. Further, not all functions are equally important, particularly in the case of contributions to services. Fortunately, choosing an ‘optimal’ level of function to link to 1 can alleviate this (e.g., if 25% of function is sufficiently high for the provision of a service, 25% or higher can be considered a ‘1’). Functions can also be upweighted or downweighted in the calculation of ^*q*^*N* so long as ∑*p*_*i*_ = 1. Choices for standardization are often best made in the context of a specific system or application and must be transparently justified.

### Correlated functions and multifunctionality

A great deal of debate in the multifunctionality literature has sprung from the issue of how to deal with correlations between functions (Bradford et al. 2014a, b, Byrnes et al. 2014b, Manning et al. 2018). As with species diversity, the rationale is that not all functions are equally different and that a metric of true multifunctionality should identify “variables that represent independent aspects of ecosystem functioning” (Manning et al. 2018). As an illustration, consider a scenario where we measure a set of functions, several of which result from a shared mechanism so that they are inextricably linked (for example, growth rate and final biomass). If we want to study the circumstances under which overall multifunctionality is maximized, the results will be disproportionately driven by circumstances maximizing this mechanism influencing the set of correlated functions; thus a naive application of a multifunctionality measure would implicitly upweight the importance of this function cluster over uncorrelated functions.

In the Ecosystem service literature, this has led to the concept of Ecosystem service bundles (Raudsepp-Hearne et al. 2010), where cluster analysis is used to identify groups (or bundles) of ecosystem services that tend to occur together. Following the same logic, Manning et al. (2018) proposed performing a cluster analysis of ecosystem functions and building a dendrogram representing the distance matrix between functions. The performance of single functions is then averaged within clusters and multifunctionality is calculated among clusters and not individual functions. This is conceptually very similar to metrics of phylogenetic and functional species diversity that construct phylogenetic trees or dendrograms based on functional similarities and measure species diversity taking into account the dendrogram structure (e.g. Faith’s PD, Allen’s H, Petchey and Gaston’s FD, Rao’s Quadratic entropy etc.). The problem is that, as discussed by Chao et al. (2014a), dendrograms are very sensitive to the type of clustering method used and the shape of a dendrogram can vary substantially between different methods (Poos et al. 2009). For the method proposed by Manning et al 2018, it is also unclear at what depth the dendrogram should be cut (i.e., what clusters are considered as independent). Fortunately, extensions of the Hill numbers framework to account for species similarity, for example in terms of traits or genetic relatedness (Chao et al. 2014a, 2019), provide a solution to the challenge of evaluating multifunctionality across correlated functions.

To account for correlation among functions in the effective multifunctionality framework, we need to incorporate several concepts from Chao et al. (2019). First, how similar are functions? For this, we need a distance matrix of some sort (Table 1). If this is based on *a priori* mechanistic information about the underlying processes driving functions (e.g., physiological or evolutionary constraints), so much the better. We often do not have such a matrix. Other options include a matrix derived from principal components or other methods of constructing distance matrices (see Manning et al. 2018). In the absence of such information, a practical choice could be to look at the correlation matrix among functions, *R*. Defining *D* = (1 - *R*)/2 would create a distance matrix where a *d*_*ij*_ value of 0 means two functions are inextricably linked (perfectly correlated) while 1 means they trade-off completely (*r*_*ij*_ = -1). This is one choice. We note that some functions could be correlated for non-biological reasons, and as such a biologically-based distance matrix might be wiser. We suggest that determining the proper way to create a distance matrix between functions for the estimation of effective diversity is an exciting area of research.

With a distance matrix in hand, we need to ask ourselves, how distinct must two functions be before we consider them completely different? This incorporates the threshold τ (Table 1) proposed in Chao et al. (2019). Using this threshold, the distance matrix is truncated such that d_ij_(τ) = min(d_ij_, τ). If, for example, τ ≤ min(d_ij_), all functions are equally distinct and we recover the results ignoring correlations. If τ = 1 or the maximum distance in the matrix, we fully incorporate information from the distance matrix with no truncation. If τ = 0.5, all negatively correlated functions are as distinct as uncorrelated functions. Chao et al. (2019) recommend using the average weighted pairwise distance between any two functions as τ

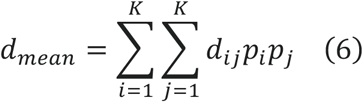

as setting τ to d_mean_ yields results that are consistent with comprehensive evaluations of all possible values of τ.

We can then use our truncated distance matrix, *d*_*ij*_(τ), to define the effective number of underlying functions, ^*q*^*N*_*u*_ as follows when we are interested in the effective number of underlying “functions”. This equation takes advantage of *d*_*ij*_(τ)/τ signifying the portion of one function that is distinct from another.

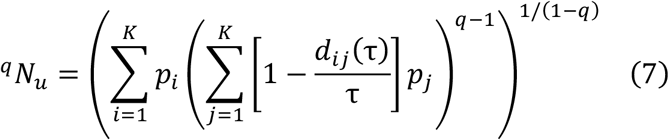

In essence, the inner part of the equation is the proportion of function *i* multiplied by the summed portion of all functions that contribute to the same underlying function. For *q* = 1, we need to use the limit, as before, so

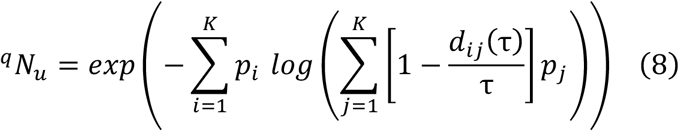

This effective number of functions can then be used to calculate multifunctionality as above. The average, *A*, does not need to be modified, because correlations do not systematically bias estimates of the mean (although uncertainty in that mean could be biased downwards by correlations). The correlations only need to be accounted for in the estimate of ^*q*^*N*.

## Discussion

With the framework above in hand, we are hopeful for a proliferation of literature investigating both new and old ideas in multifunctionality. With its ties to theoretically grounded methods of measuring species diversity and its flexibility to encompass some of the real sticking points in the field, we hope that Ecology will embrace this framework moving forward. It produces intuitive metrics - effective number of functions performing at equal levels, effective multifunction scaled by total level of performance, and standardized versions of both - as well as offering a solution to account for correlated functions. To aid in this advance, see Appendix 2 where we present a worked example in R (R Core Development Team 2008). Over the last decade and a half of its development, the multifunctionality literature has grown more slowly than it should, in no small part due to the *ad hoc* nature of the metrics we have developed. We hope that period is at an end.

Further, the metrics presented here are the foundation of a much larger framework that has seen deep exploration in the species diversity world. By embracing this framework for Multifunctionality research, we open up new vistas for ecology. Some are small - what are the consequences and best choices for τ, how we calculate distance matrices, at what scale to evaluate correlation between functions, should we incorporate changes in correlation and distance matrices as an ecological response in and of themselves, how we standardize functional measurements, and many more. Other new areas of inquiry are quite large. Can we use this framework to begin to address the problem of unmeasured functions as we deal with unmeasured species (Chao et al. 2014b)? Can we use it to think about turnover in multifunctionality across space and time as we do with beta diversity (Chao and Ricotta 2019) or think about partitioning landscape multifunctionality into different components (Jost 2007)? Particularly as we think about global change’s impacts on ecosystem services, and not just functions, at multiple scales, this framework lays out fruitful avenues for future exploration.

What does our approach mean for previous metrics? In short, existing approaches have addressed specific aspects of measuring multifunctionality, but are limited in their scope. Here we provide a metric that addresses the generalized measurement of multifunctionality that meets its definition – the “simultaneous performance of multiple functions” (Byrnes et al. 2016) – in a way not fully encompassed by previous metrics. The averaging metric comes closest to what is presented here, as it is a special case where q = 0. In the supplementary materials we provide an example using data from Duffy et al. 2003 and the *multifunc* package (v 0.9.4) in R, now including metrics in this paper, that shows some concordance between the two. The threshold approach seeks to remedy some of the drawbacks of using q = 0, and can be useful for managers that have specific targets of functional performance they need to meet. However, it is too sensitive to a number of decisions, as well as complicated to interpret (Gamfeldt and Roger 2017). Other metrics in the literature are often targeted at questions related to multifunctionality, but do not directly address a univariate measure of simultaneous function (e.g., the approach of Dooley et al. 2015 seeks to look at a multivariate response and quantify tradeoffs rather than provide a single univariate measure). Rather, we have sought to provide a metric that builds on past work while providing a robust foundation for the future of multifunctionality research.

Ultimately, we feel that the proliferation of univariate multifunctionality metrics without strong theoretical underpinning has caused a great deal of confusion about how to measure multifunctionality. We hope that this piece will provide the field of multifunctionality with a way out of its current state of division and confusion. Further, we hope it provides food for additional theory that addresses the causes and consequences of ecosystem multifunctionality, something that is currently sorely lacking but highly relevant to policy and management (e.g., the efforts of the Intergovernmental Science-Policy Platform on Biodiversity and Ecosystem Services). We have been heartened by the idea, leaving the cradle of the field of biodiversity and ecosystem function, and feel that it has the promise to provide a holistic unifying concept for anyone interested in capturing a snapshot of system dynamics in a single meaningful metric with direct ties to the beautifully developing field of diversity partitioning. Much is to be done on honing the particulars of this approach, but we feel it offers a strong theory-driven unified approach that will enable the field of Multifunctionality research to move forward swiftly.

## Supporting information

Appendices to Understandable Multifunctionality Measures Using Hill Numbers

## Acknowledgements

We thank Mark Urban and Morgan Tingley for organizing the workshop that began informal discussions between JEKB and RB leading to this manuscript. We thank Christian Alsterberg for giving reason for FR to travel to Boston where FR and JEKB realized that the three authors had independently come to the same conclusions and should collaborate instead of working in competition. It’s how science should be. We also thank funding from multiple seminar series and statistical workshops that have kept the collaboration going. This work was partially supported by the National Science Foundation as part of the PIE-LTER Program (award #1637630).

## Data Access

In addition to the appendices, see https://github.com/jebyrnes/new_multifunc_metric/ for additional code, notes, and co-author discussions related to this manuscript. The code to implement all methods here is part of the *multifunc* package in R and can be found at https://jebyrnes.github.io/multifunc/ with tutorials.

