## Appendices to Understandable Multifunctionality Measures Using Hill Numbers for "Understandable Multifunctionality Measures Using Hill Numbers"

### Appendix 1: Lower Bound of Effective Number of Functions

The minimum number of effective functions will occur at the minimum level of evenness (maximum level of dominance) for any set of functions. Conveniently, using a standardized scale for functions between 0 and 1, the maximum level any function can achieve is 1. If all functions, save one, are 0, then by definition the number of effective functions will be 1. After this, one function will be 1 and all functions save one other will be 0. As we increase $A$, this function will increase in value until it, too, is 1. At which point, as $A$ increases, a third function will begin increasing in value, etc.

Noting that $AK$, the average level of function multiplied by the total number of functions, is always just greater than the number of functions that are equal to 1, we can a quantity $G$ that is the number of functions equal to 1, and is equivalent to the floor of $AK$ (i.e., $G = \lfloor AK \rfloor$).

For any level of A, the value of the function whose value is greater than 0 but less than 1 is $AK - G$, as $AK$ is the number of functions that are 1 + the additional contribution of our non-zero other function. Therefore,

$$F_1 ... F_G = 1$$
$$F_{G+1} = AK - G$$
$$F_{G+2} ... F_K = 0$$

and therefore summing up all of the functions at the maximum dominance for a given level of $A$,

$$\sum F_i = G + AK - G + 0$$
$$\Rightarrow AK$$

We can use this to determine the lower bound of the effective number of functions. Looking at $p_i$ at maximum dominance

$$p_i = \frac{F_i}{\sum F_i}$$
$$\Rightarrow \frac{F_i}{AK}$$

We can then substitute this into equation 3.

$$^qN = \left[ \sum_{i=1}^{K} \left( \frac{F_i}{AK} \right)^q \right]^{1/(1-q)}$$

$$\Rightarrow \left[ \frac{1}{A^q \cdot K^q} \cdot \sum_{i=1}^{K} F_i^q \right]^{1/(1-q)}$$

At maximum dominance we know that

$$\sum_{i=1}^{K} F_i^q = G + (AK - G)^q$$

So that the lower bound of effective number of functions is

$$^qN_{lower}(A) = \left[ \frac{G + (AK - G)^q}{A^qK^q} \right]^{1/(1-q)}$$

For q=1, we can find the lower bound by taking the limit $\lim_{q \to 1}$, which gives us

$$\lim_{q \to 1} {}^qN_{lower}(A) = AK(AK - G)^{\frac{G}{AK} - 1}$$

In the limiting case of $AK \to G$, the expression becomes $AK$

We can write a function in R to explore this relationship between $^qN$ and $A$.

```
get_mfn_floor <- function(a, k, q){
  ak <- a*k
  g <- floor(a*k)
  if(a == 0)
    return(0)

  if(q==1){
    if(ak-g==0){ #Shannon case limit
      #AK ^ {\frac{G}{AK}}
      return(ak)
    }else{ #Shannon case

      #\lim_{q \to\ 1} ~ ^qN_{lower}(A) =  A K (A K - G)^{\frac{G}{AK} - 1}
      ret <- ak*(ak-g)^(g/ak - 1 )

    }
  } else{ #anything else where q > 1

  #\left[ \frac{G + (AK-G)^q}{A^qK^q} \right]^{1/(1-q)}
  ret <- ( (g+(ak-g)^q)/(a^q * k^q) ) ^ (1/(1-q))

  }
  return(ret)
}
```

We can use this function to then explore the shape of the relationship between A and possible values of $^qN$ which is reported in the text as figure 1.

```
library(dplyr)
library(tidyr)
library(ggplot2)
library(forcats)

n_func <- 4

n_range <- crossing(data.frame(a = seq(0.0001,1,length.out=200)),
                    data.frame(q = 1:4)) %>%
  group_by(q, a) %>%
  mutate(n_upper=n_func, n_lower=get_mfn_floor(a, n_func, q))


ggplot(n_range, aes(x=a, y=n_lower,
                          ymin = n_lower,
                          ymax = n_upper,
                          fill = fct_rev(factor(q)))) +
  geom_ribbon(alpha=0.7) +
  xlab("Average Level of Functioning") +
  ylab("Effective Number of Functions") +
  ylim(c(0,n_func)) +
  scale_fill_brewer(palette="Accent") +
  guides(fill=guide_legend("Order (q)")) +
  theme_bw(base_size = 14)
```

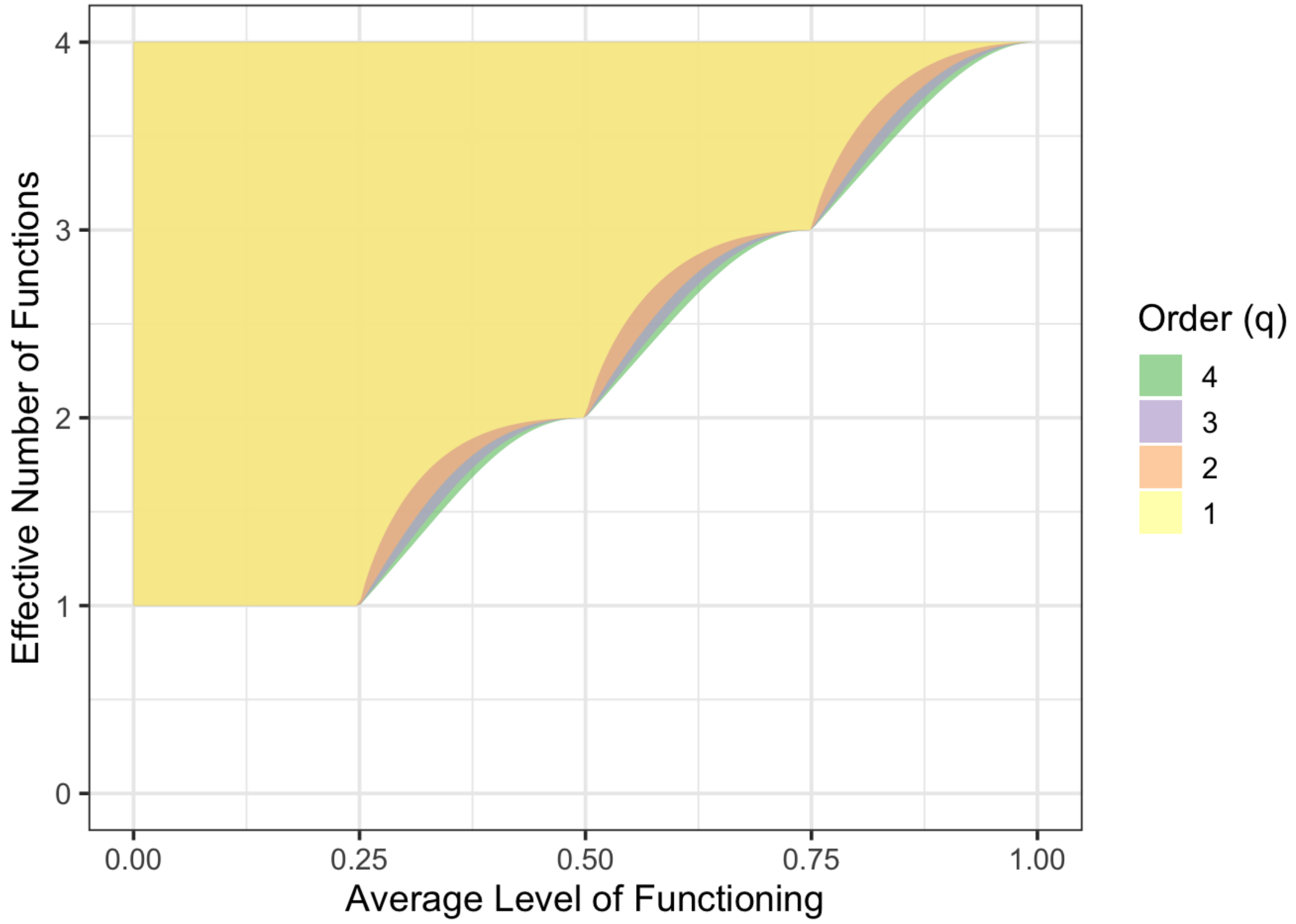

### Appendix 2 - A Worked Example with Data from Duffy 2003

To see how effective number of functions and effective multifunctionality can be used in practice, we have added functions for both into the `multifunc` package in R. To see how these metrics can be used, consider the example of Duffy et al. 2003. In this experiment, Duffy and colleagues sought to examine how biodiversity of grazers influences multiple different ecosystem functions in seagrass ecosystems. Using this framework, we standardized and reflected them as per how Duffy et al. discuss their results. We then compare average functional performance, effective multifunctionality with q = 1 and 2, and the same accommodating for correlations between functions.

### Data prep and initial calculations

First, let's load up the data, reflect two functions for which lower values mean higher levels of function.

```
library(multifunc)
library(dplyr)
library(tidyr)
library(forcats)
library(ggplot2)
theme_set(theme_bw(base_size = 14))

data("duffy_2003")
duffyAllVars <- qw(grazer_mass,wkall_chla,tot_algae_mass,
                  Zost_final_mass,sessile_invert_mass,sediment_C)

#re-normalize so that everything is on the same
#sign-scale (e.g. the maximum level of a function is the "best" function)
#and the dataset is cleaner
duffy <- duffy_2003 %>%
 dplyr::select(id, treatment,
               diversity, all_of(duffyAllVars)) %>%
 dplyr::mutate(wkall_chla = -1*wkall_chla +
                 max(wkall_chla, na.rm=T),
               tot_algae_mass = -1*tot_algae_mass +
                 max(tot_algae_mass, na.rm=T))
```

We can then plot the functions and produce a figure similar to the one in the original manuscript - although slightly different given the reflected functions.

```
# Plot functions we want
duffy %>%
  select(treatment, diversity, all_of(duffyAllVars)) %>%
  pivot_longer(all_of(duffyAllVars)) %>%
  ggplot(aes(x = diversity, y = value,
             color = treatment)) +
  stat_summary(fun.data = mean_se) +
  stat_smooth(method = "lm", aes(group = name)) +
  facet_wrap(vars(name), scale = "free_y") +
  scale_color_brewer(palette = "Set3") +
  labs(y = "Level of Function", x = "Species Richness",
       color = "Treatment")
```

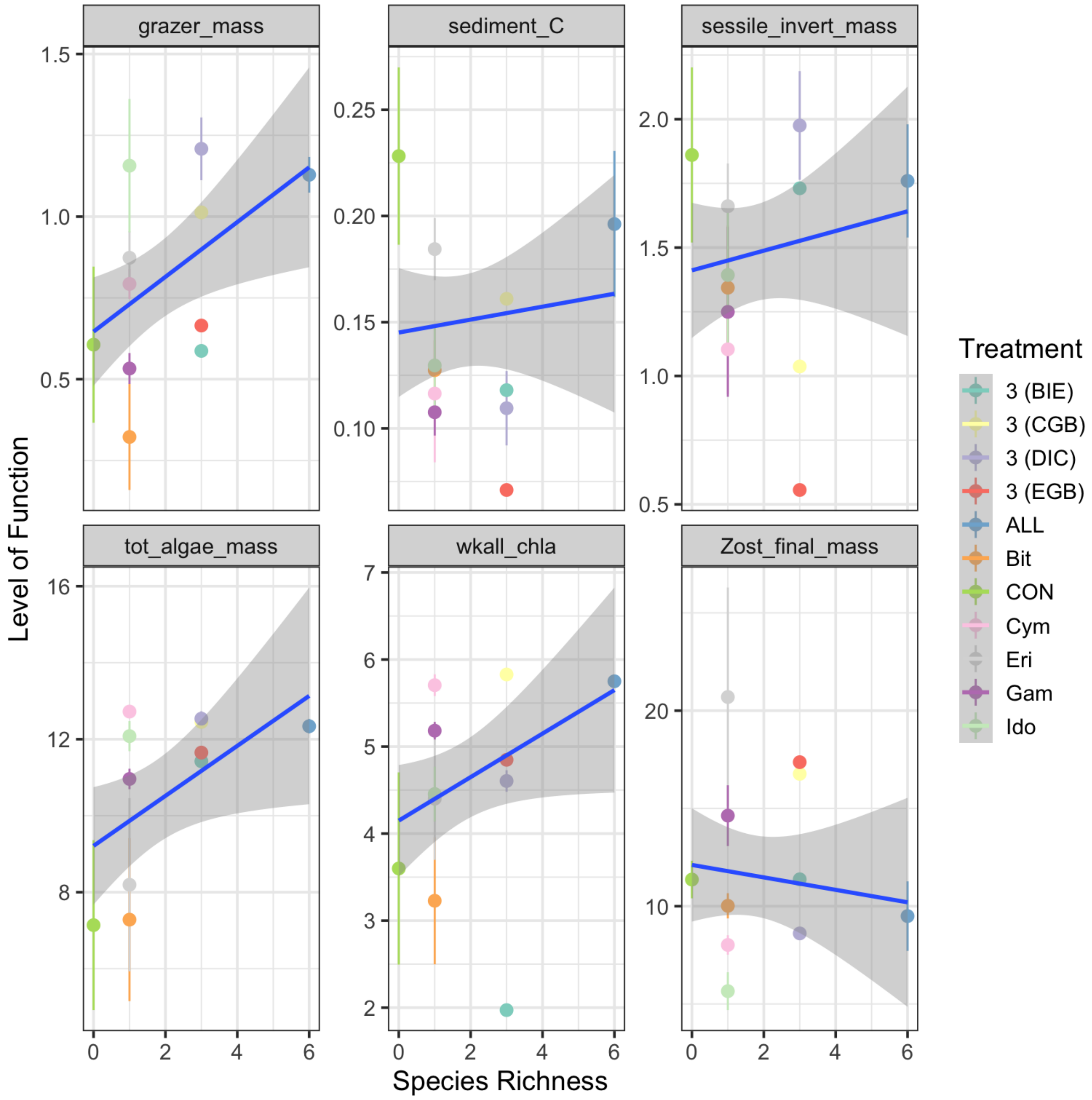

From this, we can standardize functions and calculate A, the mean multifunctionality.

```
#first, mean multifunctionality
duffy <- duffy %>%
 cbind(getStdAndMeanFunctions(duffy, duffyAllVars)) %>%
 dplyr::rename(richness=diversity)



duffy <-duffy %>%
  getFuncsMaxed(duffyAllVars,
                threshmin=0.8, threshmax=0.8)
```

We can calculate effective number of functions with `multifunc::eff_num_func()` , which has arguments for different levels of q, standardizing by number of functions, and the use of distance matrices and tau. Let's calculate the effective number of functions for q = 1 and q = 2. We can then multiple both by the average to get effective multifunctionality.

```
duffyAllVars.std <- paste0(duffyAllVars, ".std")

#now effective number of functions
duffy <- duffy %>%
  mutate(n_eff_func_1 = eff_num_func(., duffyAllVars.std, q = 1),
         n_eff_func_2 = eff_num_func(., duffyAllVars.std, q = 2),

         mf_eff_1 = n_eff_func_1 * meanFunction,
         mf_eff_2 = n_eff_func_2 * meanFunction
         )
```

### Comparison of approaches

With these metrics in hand, how different do either effective number of functions or effective multifunctionality look from previous approaches?

```
duffy %>%
  select(id, treatment, richness,
         meanFunction, funcMaxed,
         n_eff_func_1, n_eff_func_2,
         mf_eff_1, mf_eff_2) %>%
  pivot_longer(cols = c(meanFunction:mf_eff_2)) %>%
  mutate(name = fct_inorder(name)) %>%

  #now a plot
  ggplot(aes(x = richness, y = value,
             color = treatment)) +
  stat_summary(fun.data = mean_se) +
  stat_smooth(method = "lm", color = "black") +
  facet_wrap(vars(name), ncol = 2, scale = "free_y") +
  scale_color_brewer(palette = "Set3")
```

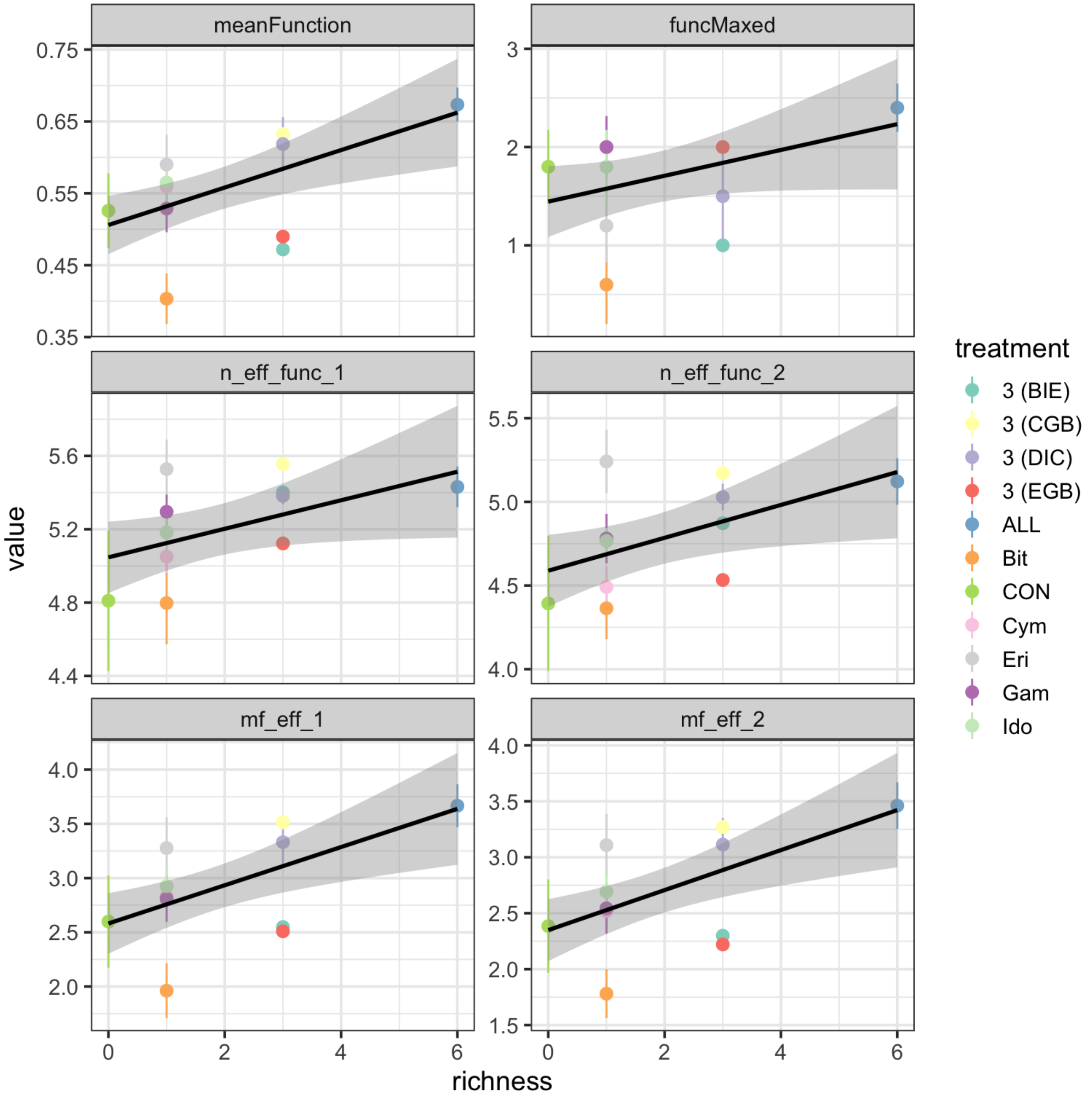


Comparing across all metrics, they produce similar results - as one would hope. Note, however, that the drop-down in average for some of the intermediate diversity treatments is not as strong for effective number of functions. While we see it's influence again in effective multifunctionality, note that some of the treatments have actually reversed in rank order. This is even more apparent when comparing to number of functions > 0.8, which is also flatter. These observations that the story described from the effective multifunctionality approaches are similar - at least in this data set - to those told by the averaging approach. However, they are subtly different in some key elements, which could have consequences for interpretation, and they have a much more solid grounding in Ecological theory.

Further, the effective multifunctionality results broadly reproduce the qualitative conclusions of Duffy et al. (2003). They reveal, however, that in lower diversity assemblages, there is less of a drop in effective number of functions. The reason for the strong Biodiversity-Multifunctionality results derive from three observations. First, some lower diversity treatments have low average levels of function despite diversity having more mild impact on evenness of function. Second, the control treatment, while maintaining a high level of function, has a much lower effective number of functions. Thus, the skew in functional dominance in the control drives down effective multifunctionality in this treatment at 0 richness. Third, some monocultures have both low mean and low effective number of functions, further strengthening the effective multifunctionality results beyond what is seen in either the mean, threshold, or number of effective functions results alone.

### Adjusting for correlation between functions

We can also incorporate correlation between functions to calculate effective number of functions controlling for some functions being driven by the same underlying unmeasured function - this producing lower scores for effective number of functions than would appear from our initial approach.

To begin with, we will need a distance matrix based on correlations. For that, we have `multifunc::cor_dist()` which calculates a correlation matrix, subtracts it from 1 and divides by two to make a distance matrix.

```
D <- cor_dist(duffy %>% select(all_of(duffyAllVars.std)))
```

With the approach outlined for effective multifunctionality, we need to also consider a cutoff in similarity, $\tau$. As a default choice for functions, we use $d_{mean}$, the average of the matrix weighted by the total frequency of function. To see what $d_{mean}$ would be, we can use `multifunc::dmean()` and then either use that in future functions, or recognize that functions default to calculating $d_{mean}$. Other choices could be $d_{max}$ or $d_{min}$ - the minimum non-zero value of the distance matrix which can be calculated with `multfunc::dmin()` in the same way we use `dmean()` below.

```
tau <- dmean(duffy %>% select(all_of(duffyAllVars.std)),
             D)

tau
#> [1] 0.3860855
```

We can then get the effective multifunctionality for q = 1 and q = 2 adjusting for correlation between species. Here we will use `multifunc::getMF_eff()` so that we do not have to calculate effective number of functions and multiply by the mean. We can though, if we would like, using `n_eff_func()` as before with new arguments as below.

```
duffy <- duffy %>%
  mutate(mf_eff_1_cor = getMF_eff(., duffyAllVars, q = 1,
                                  D = D, tau = tau),
         mf_eff_2_cor = getMF_eff(., duffyAllVars, q = 2,
                                  D = D, tau = tau)
         )
```

Let’s then evaluate the consequences of incorporating correlation.

```
duffy %>%
  select(id, treatment, richness,
         mf_eff_1, mf_eff_2,
         mf_eff_1_cor, mf_eff_2_cor) %>%
  pivot_longer(cols = c(mf_eff_1:mf_eff_2_cor)) %>%
  mutate(name = fct_inorder(name)) %>%

  #now a plot
  ggplot(aes(x = richness, y = value,
             color = treatment)) +
  stat_summary(fun.data = mean_se) +
  stat_smooth(method = "lm", color = "black") +
  facet_wrap(vars(name), ncol = 2) +
  scale_color_brewer(palette = "Set3")
```

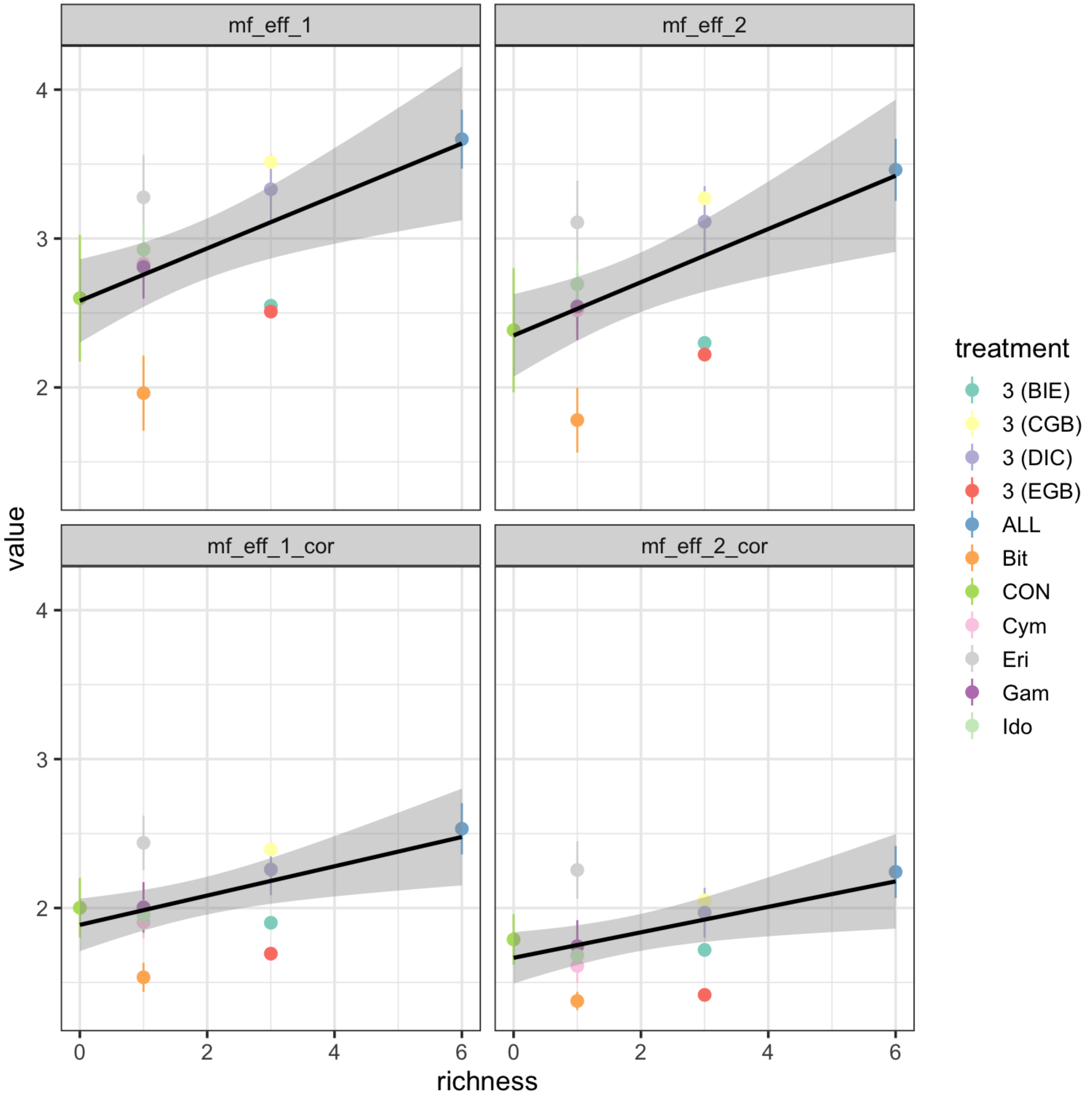


Qualitatively, the results do not appear to change very much, although values incorporating correlations are lower, as expected. However, as fewer unique underlying "functions" are being affected, the results are somewhat weaker in slope.

```
# ------------------------------------------------------
# Appendix 3: Functions added to the multifunc package
# hosted at https://jebyrnes.github.io/multifunc/ for
# the calculation of effective multifunctionality from
# Byrnes, Roger, and Bagchi's Understandable
# Multifunctionality Measures Using Hill Numbers
# ------------------------------------------------------

#' @title getMF_eff
#' @description A multifunctionality index rooted in Hill numbers.
#' \code{getMF_eff} get multifunctionality index defined by function and
effective number of functions
#'
#' @details Takes a data frame, variable names, a standardizing function,
whether we want
#' an index standardized by number of functions or not, an order of Hill
number for our
#' effective number of functions as well as a dissimilarity matrix (if
desired) and value
#' for a dissimilarity cutoff (defaults to the average dissimilarity). It
then calculates
#' both the average standardized function in each plot and the effective
number of
#' functions and returns their product as a measure of effective
multifunctionality.
#'
#' @author Jarrett Byrnes.
#' @param data A data frame with functions in columns and rows as replicates
as well as other information.
#' @param vars Name of function variables
#' @param standardized Use standardized number of functions (scaled by total
#' number of functions, so between 0-1), or just raw effective number of
#' functions for calculation. Defaults to \code{FALSE}.
#' @param standardize_function A function to standardize each individual
#' function to the same scale, such as \code{standardizeUnitScale} or
#' \code{standardizeZScore}
#' @param q  Order of the diversity measure. Defaults to the
#' Shannon case where q = 1. For Simpson, q=2.
#' @param D A distance matrix describing dissimilarity between functions.
Defaults
#' to NULL, and the index is calculated assuming all functions are different.
If
#' it is not null, it must be a symmetric matrix with dimensions matching the
#' number of functions listed in \code{vars}.
#' @param tau A cutoff for degree of dissimilarity under which functions are
considered
#' to be different. If tau is the minimum non-zero value of D, all functions
are different.
#' if tau is the maximum value of D are greater, all functions are considered
the same.
#' @references
#'
#' Chao, A., Chiu, C.-H., Villéger, S., Sun, I.-F., Thorn, S., Lin, Y.-C.,
#' Chiang, J.-M. and Sherwin, W. B. 2019. An attribute-diversity approach to
```

```
#' functional diversity, functional beta diversity, and related
(dis)similarity
#' measures. Ecological Monographs. 89: e01343.
#'
#' Jost, L. 2006. Entropy and diversity. Oikos 113(2): 363-375.
#'
#' Hill, M. 1973. Diversity and evenness: A unifying notation and its
#' consequences. Ecology 54: 427-432.
#'
#' @export
#' @return Returns a vector of effective or standardized effective
multifunctionality.
#'

getMF_eff <- function(data, vars, q = 1,
                      standardized = FALSE,
                      standardize_function = standardizeUnitScale,
                      D = NULL, tau = NULL) {

  # get standardized functions
  std_funcs <- getStdAndMeanFunctions(data, vars, standardize_function)
  funcs <- std_funcs[, -ncol(std_funcs)]

  # get the average functional level
  mf_a <- std_funcs$meanFunction

  # get evenness
  eff_func <- eff_num_func(
    dat = funcs,
    vars = names(funcs),
    q = q,
    standardized = standardized,
    D = D, tau = tau
  )

  # mf = average * eff num functions
  mf_a * eff_func
}


#' @title eff_num_func
#'
#' @description Calculate the effective number of functions for rows in a
dataset
#'
#' @param dat A data frame with functions in columns and rows as replicates
as well as other information.
#' @param vars Column names of function variables
#' @param standardized Use standardized number of functions (scaled by total
#' number of functions, so between 0-1), or just raw effective number of
#' functions for calculation. Defaults to \code{FALSE}.
#' @param q  Order of the diversity measure. Defaults to the
#' Shannon case where q = 1. For Simpson, q=2.
```

```
#' @param D A distance matrix describing dissimilarity between functions.
Defaults
#' to NULL, and the index is calculated assuming all functions are different.
If
#' it is not null, it must be a symmetric matrix with dimensions matching the
#' number of functions listed in \code{vars}.
#' @param tau A cutoff for degree of dissimilarity under which functions are
considered
#' to be different. If tau is the minimum non-zero value of D, all functions
are different.
#' if tau is the maximum value of D are greater, all functions are considered
the same.
#'
##' @details Takes a data frame, variable names,  whether we want
#' an index standardized by number of functions or not, an order of Hill
number for our
#' effective number of functions as well as a dissimilarity matrix (if
desired) and value
#' for a dissimilarity cutoff (defaults to the average dissimilarity). It
then calculates
#' and returns the effective number of functions using the appropriate
method. See Chao
#' et al. 2019 for more.
#'
#'
#' @references
#'
#' Chao, A., Chiu, C.-H., Villéger, S., Sun, I.-F., Thorn, S., Lin, Y.-C.,
#' Chiang, J.-M. and Sherwin, W. B. 2019. An attribute-diversity approach to
#' functional diversity, functional beta diversity, and related
(dis)similarity
#' measures. Ecological Monographs. 89: e01343.
#'
#' Jost, L. 2006. Entropy and diversity. Oikos 113(2): 363-375.
#'
#' Hill, M. 1973. Diversity and evenness: A unifying notation and its
#' consequences. Ecology 54: 427-432.
#'
#'
#' @return Returns a vector of effective or standardized effective number of
functions
#' @export
#'

eff_num_func <- function(dat, vars, q = 1,
                         standardized = FALSE,
                         D = NULL, tau = NULL) {

  # get the variable data frame
  adf_raw <- dat[, which(names(dat) %in% vars)]

  # calculate tau if needed
  if (!is.null(D) & is.null(tau)) tau <- dmean(adf_raw, D)
```

```
  # get relative levels of each function to all others per row
  adf_freq <- adf_raw / rowSums(adf_raw)

  # proportion of community
  if (is.null(D)) {
    ret <- eff_num_func_no_d(adf_freq, q = q)
  } else {
    ret <- eff_num_func_d(adf_freq, q = q, D = D, tau = tau)
  }

  # standardize if needed
  if (standardized) {
    ret <- ret / length(vars)
  }

  return(ret)
}

# effective # functions not adjusting for correlation
#' @title eff_num_func_no_d
#'
#' @param adf_freq A data frame of functional "frequencies" - i.e. f_i/
sum(f_i)
#' @param q Order of hill number used for index. Defaults to q=1, as in
Shannon Diversity
#' @details Takes a data frame, with functions standardized against total
level of function
#' in their replicate as columns and replicates as rows. Returns the
effective number of functions using the appropriate method. See Chao
#' et al. 2019 or Jost 2006 for details. Does not adjust for correlation
between functions.
#' @references
#'
#' Chao, A., Chiu, C.-H., Villéger, S., Sun, I.-F., Thorn, S., Lin, Y.-C.,
#' Chiang, J.-M. and Sherwin, W. B. 2019. An attribute-diversity approach to
#' functional diversity, functional beta diversity, and related
(dis)similarity
#' measures. Ecological Monographs. 89: e01343.
#'
#' Jost, L. 2006. Entropy and diversity. Oikos 113(2): 363-375.
#'
#' Hill, M. 1973. Diversity and evenness: A unifying notation and its
#' consequences. Ecology 54: 427-432.
#'
#' @return Returns a verctor of effective or standardized effective number of
functions
#' @export
#'

eff_num_func_no_d <- function(adf_freq, q = 1) {
  if (q == 1) {
    summand <- adf_freq * log(adf_freq)
    # deal with 0 log 0 = 0
    zero_idx <- which(is.nan(as.matrix(summand)), arr.ind = TRUE)
```

```
    if (length(zero_idx) > 0) {
      summand[which(is.nan(as.matrix(summand)), arr.ind = TRUE)] <- 0
    }

    exp(-rowSums(summand))
  } else {
    rowSums(adf_freq^q)^(1 / (1 - q))
  }
}

# effective # functions adjusting for correlation
#' eff_num_func_d
#'
#' @param adf_freq A data frame of functional "frequencies" - i.e. f_i/
sum(f_i)
#' @param q Order of hill number used for index. Defaults to q=1, as in
Shannon Diversity
#' @param D A distance matrix describing dissimilarity between functions.
#' @param tau A cutoff for degree of dissimilarity under which functions are
considered
#' to be different. If tau is the minimum non-zero value of D, all functions
are different.
#' if tau is the maximum value of D are greater, all functions are considered
the same.
#'
#' @references
#'
#' Chao, A., Chiu, C.-H., Villéger, S., Sun, I.-F., Thorn, S., Lin, Y.-C.,
#' Chiang, J.-M. and Sherwin, W. B. 2019. An attribute-diversity approach to
#' functional diversity, functional beta diversity, and related
(dis)similarity
#' measures. Ecological Monographs. 89: e01343.
#'
#'
#' @return A vector of effective number of functions
#' @export
#'

eff_num_func_d <- function(adf_freq, q = 1, D, tau = NULL) {
  # make sure we can do this
  check_d_and_adf_errors(adf_freq, D)

  # should have supplied one! cannot calculate
  if (is.null(tau)) stop("You need to choose a value for tau.")
  if (tau == 0) {
    warning("Tau cannot be 0, as this would lead to division by 0. Changing
to the lowest value > 0")
    tau <- min(D[D != 0])
  }
  # make sure it's a matrix!
  D <- as.matrix(D)

  # truncate by tau
  D[which(D > tau, arr.ind = TRUE)] <- tau
```

```
  # iterate over rows
  apply(adf_freq, 1, eff_num_func_d_onerow, D = D, tau = tau, q = q)
}


#' eff_num_func_d_onerow
#'
#' @param arow_freq One replicate sample of different functions (a single
numeric vector)
#' @param q Order of hill number used for index. Defaults to q=1, as in
Shannon Diversity
#' @param D A distance matrix describing dissimilarity between functions.
#' @param tau A cutoff for degree of dissimilarity under which functions are
considered
#' to be different. If tau is the minimum non-zero value of D, all functions
are different.
#' if tau is the maximum value of D are greater, all functions are considered
the same.
#' @references
#'
#' Chao, A., Chiu, C.-H., Villéger, S., Sun, I.-F., Thorn, S., Lin, Y.-C.,
#' Chiang, J.-M. and Sherwin, W. B. 2019. An attribute-diversity approach to
#' functional diversity, functional beta diversity, and related
(dis)similarity
#' measures. Ecological Monographs. 89: e01343.
#'
#'
#' @return A single value of effective number of functions
#' @export
#'

eff_num_func_d_onerow <- function(arow_freq, D, tau, q) {
  arow_freq <- as.numeric(arow_freq)
  a <- as.vector((1 - D / tau) %*% arow_freq) # from Chao et al 2019 code

  if (q == 1) {
    exp(-1 * sum(arow_freq * log(a)))
  } else {
    sum(arow_freq * a^(q - 1))^(1 / (1 - q))
  }
}


#' @title dmean
#' @description Calculates the average distance between functions for one or
an entire
#' assemblage of replicates
#'
#' @param adf_raw A data frame frame with functions in columns and rows as
replicates
#' @param D A distance matrix describing dissimilarity between functions.
#' @references
#'
```

```
#' Chao, A., Chiu, C.-H., Villéger, S., Sun, I.-F., Thorn, S., Lin, Y.-C.,
#' Chiang, J.-M. and Sherwin, W. B. 2019. An attribute-diversity approach to
#' functional diversity, functional beta diversity, and related
(dis)similarity
#' measures. Ecological Monographs. 89: e01343.
#'
#' @return Single numeric of weighted average of distance matrix
#' @export
#'

dmean <- function(adf_raw, D) {
  # make sure we can do this
  check_d_and_adf_errors(adf_raw, D)

  # sum ( (tmp %*% t(tmp) ) * dij)
  p_vec <- colSums(adf_raw) / sum(adf_raw)

  # Q <- 0
  #
  # for(i in 1:ncol(D)){
  #   for(j in 1:ncol(D)){
  #     Q <- Q + D[i,j]*p_vec[i]*p_vec[j]
  #   }
  # }

  # from Chao et al. 2019 code - matches above
  sum(p_vec %*% t(p_vec) * D)
}


#' dmin
#'
#' @description Calculates the minimum non-zero value of a distance matrix
#'
#' @param D A distance matrix describing dissimilarity between functions.
#'
#' @return A numeric
#' @export
#'

dmin <- function(D) {
  min(D[D != 0])
}

#' @title cor_dist
#' @description Takes a data frame of functions and calculates the
correlation-based
#' distance between functions.
#'
#' @param adf A \code{data.frame} or \code{matrix} of functions
#'
#' @return A matrix
#' @export
#'
```

```
cor_dist <- function(adf) {
  if (sum(is.na(adf)) > 0) stop("Some function values are NA. Cannot compute
a distance matrix.")
  cmat <- stats::cor(adf)
  D <- (1 - cmat) / 2
  D
}

## Error checks for using distance-matrix based effective multifunctionality
## that could be needed for multiple different functions
check_d_and_adf_errors <- function(adf, D) {
  # error checks
  if (sum(is.na(adf)) > 0) stop("Some function values are NA. Cannot compute
multifunctionality for those rows. Please filter them out before running.")

  if (nrow(D) != ncol(D)) {
    stop("Distance matrix is not square")
  }

  if (!isSymmetric(D)) {
    stop("Distance matrix is not symmetric")
  }

  if (nrow(D) != ncol(adf)) {
    stop("Distance matrix does not have same number of rows as number of
variables")
  }

  if (ncol(D) != ncol(adf)) {
    stop("Distance matrix does not have same number of columns as number of
variables")
  }
}
```
